# The *openCARP* Simulation Environment for Cardiac Electrophysiology

**DOI:** 10.1101/2021.03.01.433036

**Authors:** Gernot Plank, Axel Loewe, Aurel Neic, Christoph Augustin, Yung-Lin Huang, Matthias A. F. Gsell, Elias Karabelas, Mark Nothstein, Anton J. Prassl, Jorge Sánchez, Gunnar Seemann, Edward J. Vigmond

## Abstract

**Background and Objective:** Cardiac electrophysiology is a medical specialty with a long and rich tradition of computational modeling. Nevertheless, no community standard for cardiac electrophysiology simulation software has evolved yet. Here, we present the *openCARP* simulation environment as one solution that could foster the needs of large parts of this community.

**Methods and Results:** *openCARP* and the Python-based *carputils* framework allow developing and sharing simulation pipelines which automate *in silico* experiments including all modeling and simulation steps to increase reproducibility and productivity. The continuously expanding *openCARP* user community is supported by tailored infrastructure. Documentation and training material facilitate access to this complementary research tool for new users. After a brief historic review, this paper summarizes requirements for a high-usability electrophysiology simulator and describes how *openCARP* fulfills them. We introduce the *openCARP* modeling workflow in a multi-scale example of atrial fibrillation simulations on single cell, tissue, organ and body level and finally outline future development potential.

**Conclusion:** As an open simulator, *openCARP* can advance the computational cardiac electrophysiology field by making state-of-the-art simulations accessible. In combination with the *carputils* framework, it offers a tailored software solution for the scientific community and contributes towards increasing use, transparency, standardization and reproducibility of *in silico* experiments.

## 1 Introduction

Computational modeling and simulation of cardiac electrophysiology (CEP) has emerged in the last decades and is now playing a pivotal role in basic cardiology research [1]. It also shows high promise as a clinical research tool, for device and drug development [2] and even as a complementary clinical modality, aiding in diagnosis, therapy stratification and planning in future precision cardiology [3]. A key motivation driving CEP model development is the unique ability of providing a mechanistic framework for integrating disparate experimental or clinical data gathered in *in vivo, in vitro* or *ex vivo* and subjecting these to thorough quantitative analysis in a matching *in silico* setting. Such *in silico* CEP models allow study of complex cause-effect relationships at a level of quantitative accuracy and biophysical detail beyond what is feasible today with any other research modality. Advanced *in silico* CEP models facilitate the observation of almost any quantity of interest at high spatio-temporal resolutions at scales ranging from cellular to organ, including studies on human hearts, without being limited by ethical constraints. These advantages have led to a marked increase in modeling-based or modeling-augmented publications in cardiology since the early 2000s (Figure 1) with a wide variety of different software implementations, e.g. illustrated in [4–8] or a joint verification benchmark effort of the CEP community [9].

**Figure 1:**
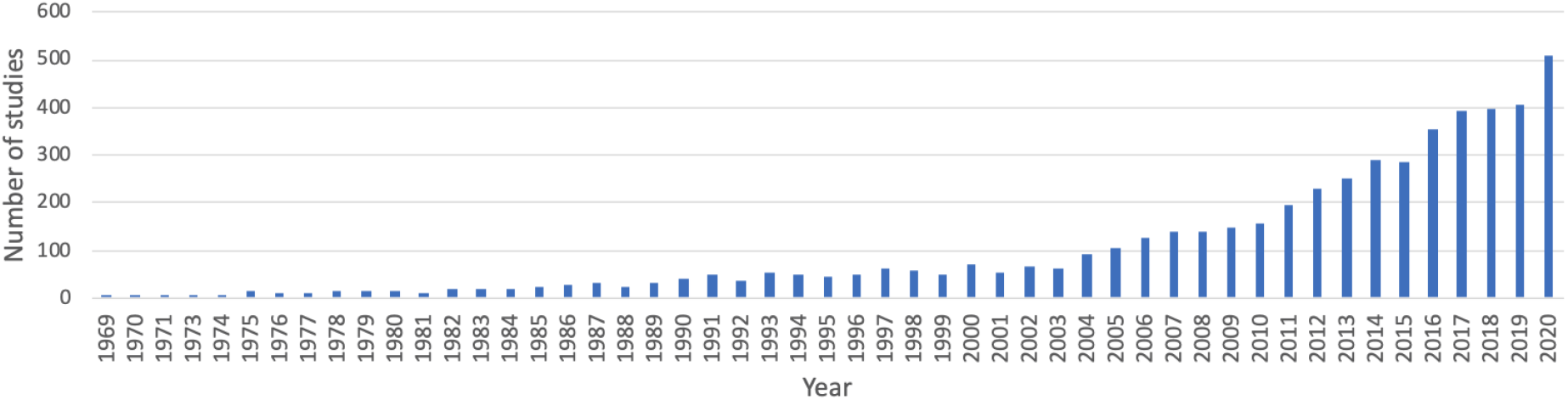
PubMed-listed studies with “(atria* OR ventric* OR cardi*) AND (“computer model” OR “mathematical model” OR “computational model” OR “in silico”)” in title or abstract.

Today, simulations of cardiac function in anatomically accurate and biophysically detailed *in silico* models have become feasible. Key factors hampering a further adoption of *in silico* CEP models in advanced application scenarios in basic research, industrial device development and clinical decision making are the limited access to cutting edge simulation technology, and the inherent complexities involved in using such sophisticated tools. These factors render setting up and performing advanced simulation studies a challenging endeavor. In this paper, we present *openCARP*, a CEP modeling environment fully open for academic use that aims to lift these limitations to increase accessibility and, thus, boost adoption of *in silico* analysis of CEP. To cope with the broad range of demands in terms of efficiency, flexibility and usability, the *openCARP* modeling environment comprises two major software components: the actual *openCARP* simulator and an open source Python framework, referred to as *carputils*, for describing *in silico* experiments. *carputils* facilitates building and sharing multiscale workflows by standardization of parameterization, execution, and archiving of simulations, to increase reproducibility and enhance robustness and reliability of complex CEP studies. In general, *carputils* can be adapted to support other CEP simulators as well. Infrastructure supporting interaction combined with extensive tailored documentation and regular user meetings provide the basis for fostering a vibrant user community.

This paper first gives a brief historic overview of the field of CEP simulation software, from the early pioneering days to the current situation that motivated the development of the *openCARP* simulator and *carputils*. Then, we review requirements for a CEP simulation environment to accommodate the needs of a wide user base. We introduce the *openCARP* CEP simulator and the *carputils* Python framework, together with pre- and post-processing components. The use of these tools is described along a typical workflow for setting up a state-of-the-art *in silico* simulation study, that spans from single cell to organ level, covering a wide range of user needs in the CEP modeling and simulation field.

### 1.1 Historical CEP Software Development

It was not before the mid-1980s that computational models for studying bioelectrical activity at the tissue level emerged [10, 11]. Unlike modeling work focused on vascular mechanics or hemodynamics, for which – owing to close similarities to applications of industrial relevance – commercial software became available early on, this was not the case for CEP modeling. In the absence of such software, academia had to develop CEP modeling solutions, which has remained that way up until today. While commercially available multiphysics simulators may have CEP modules, they are limited in terms of speed and capability [12]. In general, these do not meet the demands of state-of-the-art CEP studies which have undergone a marked transformation over the past decade, leading to a dramatic increase in complexity [13, 14].

As a consequence, the vast majority of published CEP modeling studies have relied on academic in-house codes [15–23]. These software packages have been largely developed by individual laboratories as side projects in support of their applied research work, typically focused on understanding mechanisms underlying the formation and maintenance of arrhythmias [24, 25] and their therapies [26–28]. The direction of software development has been largely steered by the needs of funded projects. Often, this led to *ad-hoc* development processes yielding research prototypes but no sustainable software products for long-term use, rather than a roadmap-based development targeting longer term strategic goals. Moreover, particularly during the pioneering years, the modeling infrastructure encoded the specific expertise of individual labs. Sharing of software or models across labs was uncommon. Research on CEP resulting in discovery has been traditionally regarded as being of higher academic merit than research on building the enabling CEP modeling methodology, which requires a similar amount of human resources. Thus, modeling has been significantly less well funded, which has opened a gap between the often highly ambitious scientific goals and the available technical capabilities [29].

For at least the first three decades of CEP modeling, the single lab paradigm was the prevailing approach in the development of CEP modeling infrastructure. An important factor rendering this approach viable was the simple nature of CEP simulations during these early years, making it feasible for a single graduate student to develop and write code for the problem at hand. Models of cellular dynamics were relatively simple [30–32], and geometric representations were anatomical abstractions, comprising 1D strands, 2D sheets or, later in the late nineties when computational resources became more powerful, 3D slab or wedge preparations. These regular domains lent themselves quite naturally to simpler spatial discretization techniques, with the finite difference method (FDM) appropriate for the vast majority of studies. Numerical techniques were less well developed and much simpler to implement, or generic implementations made available as numerical libraries were integrated.

### 1.2 Current State of CEP Modeling

Over the past decade though, the complexity of CEP simulations has increased exponentially as the result of several factors:

i. faster, and higher resolution tomographic imaging modalities
ii. an explosion of biological knowledge leading to more components in cell models, as well as more complicated formal descriptions of these components and interactions between them
iii. more elaborate numerical methods which improve performance at the cost of increased complexity
iv. evolving computer architectures which demand specific data layouts and algorithms to fully exploit the resources

Complexity has increased beyond the capabilities of a single lab development paradigm.

State-of-the-art modeling studies use unstructured, high resolution, image-based tomographic reconstructions to reflect individual cardiac anatomies with high geometric fidelity and avoid spurious boundary artefacts introduced by jagged surfaces of Cartesian grids. Unstructured tesselation has prompted more sophisticated discretization techniques, such as the finite element method (FEM) [33] or the finite volume method (FVM) [34], which are substantially more costly to implement and time-consuming to develop a profound understanding of. CEP is studied at the organ scale in biventricular or biatrial models and recently models representing all chambers of the heart are beginning to be used [35].

Such four chamber models account for anisotropic tissue properties and discrete conduction pathways in all chambers, the specialized cardiac conduction system comprising sinus node [36–38], atrio-ventricular node [39] and the His-Purkinje system [40]. Pathologies altering conduction velocities or pathways such as ischemia [41], infarct scars and surrounding transitional tissue [42], or various types of fibrotic lesions [43–45] are considered at increasing levels of physiological [46] and structural detail [47–50].

Along with the structural heterogeneity, a large number of cellular models are needed to account for functional heterogeneity across different regions of the heart, often with subtle variations within regions to account for gradients in protein expression that alter action potential (AP) morphology. The *CellML* markup language, along with its repository (cellml.org), was created to aid in the standardization and dissemination of these models [51]. Beyond the numerous issues in developing the software, major challenges in building *in silico* experiments have to be addressed. Sophisticated workflows have been conceived to facilitate automated translation of segmented image data sets into discrete meshes suitable for simulations [52–54]. These workflows are constantly refined to render the production of increasingly larger numbers of individualized models, often referred to as *virtual cohorts*, feasible [35]. In the process of functional personalization of such models, tuning parameters have to be identified to achieve a close fit with observed data. Experimental or clinical protocols are mimicked *in silico* to match *in vivo* experiments or clinical conditions ever more closely. These procedures typically require simulating prolonged periods to ensure convergence to a limit cycle, and the number of simulations executed within optimization loops tend to be high. Overall, these factors, in combination with the numerical demands imposed by CEP – the fast upstroke of the action potential translates into depolarization wave fronts of limited spatial extent (<1 mm) – render the execution of modeling studies a challenging endeavour. Such studies can only be executed with highly efficient, robust and versatile CEP simulation tools. The costs in terms of personnel necessary to develop such tools from scratch is daunting. For instance, the cost of development of *CARP*, the proprietary predecessor of *openCARP*, is estimated to be around 50 person-years, not including any of the pre- and post-processing codes necessary for building the workflows.

### 1.3 Towards a Common Software

In other fields, certain software has been adopted by a critical mass of the community and thus have become the *de facto* standard for the entire field. Examples of this are *Neuron* for neural simulation [55], and *GAMESS* for chemistry [56]. While the need for standardized, open software has been recognized before within the CEP modeling community, no software has achieved such a status. Preceding initiatives in this vein include *openCMISS* [18] and *Chaste* [57], but their success has been mixed so far, as their rate of adoption has been rather marginal. Reasons are multifactorial: For groups developing numerical methods, there is limited demand as they have the capability to build custom tailored frameworks themselves [23, 58–62] that facilitate the implementation of disruptive changes at any time, without the frictional losses involved in community projects where changes have to be agreed upon by various stakeholders. This is in stark contrast to the demands within the applied CEP modeling community. A broad web-based survey conducted by us in 2017 showed that for a convergence in the CEP field, simulation environments are needed which meet most or, ideally, all of the following requirements:

- **Feature completeness**, i.e. a simulator should have the features to be able to replicate a large share of published modeling studies using other software [57, 63]. It must be able to perform the functions of any software it is replacing, as well as offer new functionality.
- The simulator needs to be made **available** under a license that allows unrestricted academic use [64].
- A **streamlined installation** process for all popular deployment targets without the need to deal with intricate technical challenges of compilation and software dependencies [64].
- An intuitive and flexible **user interface** that exposes input parameters in an intuitive way while accommodating a wide range of experimental conditions and protocols. Consistent interfaces are the basis for a positive and intuitive user experience [65].
- Comprehensive **user documentation** for all tools, combined with **extensive training material**, such as tutorials for all simulator features to get new users started within a reasonable time, and a tractable learning curve [64].
- The simulation environment should provide streamlined, **standardized workflows** to share, reproduce, and archive *in silico* experiments [66]. It should be easy to export results, together with all relevant input data, in one bundle. A high degree of standardization also facilitates the sharing of expertise between groups.
- A vibrant **user and developer community** is key for a sustainable research software [67]. These communities should be supported by interactive platforms, spur development (feature requests, bug reports), and interact via webcasts and user meetings [64].
- The simulator should be **computationally efficient** on all common hardware platforms ranging from local workstations to national high performance computing facilities [68].
- **Code documentation** with a thorough description of classes, methods and interfaces, as well as the overall software architecture, lowers the barrier for users becoming developers and contributing to the project. This documentation, in combination with a modular and extensible architecture, is the basis for sustainable research software [69].
- Import from (e.g. *CellML* [51] for cellular models) and export to (e.g. VTK [70]) common data formats increases **interoperability**. Easy integration in existing pre- and post-processing workflows attracts a wider range of users. The version of the simulator code used to generate a result should be clearly identified and accessible via a persistent and citable identifier.
- A high level of trust in the correctness of the software implementation is instrumental. The functionality of the simulator needs to be verified and constantly controlled by **quality assurance** measures. Before entering the master branch, new code should be reviewed by maintainers. Test cases should cover all use cases and be executed for each build before deployment [64]. The necessary trust is best established in a positive feedback loop wherein a critical mass of CEP modeling labs adopts and successfully uses the software for a broad range of modeling applications over prolonged periods.

## 2 Methods and Results

The main objective of the *openCARP* initiative is to provide and establish simulation software within the CEP modeling community which fulfills the criteria outlined above. To better meet these user requirements, the *openCARP* modeling environment is organized as comprising two major software bundles, the actual code for executing simulations, i.e., the *openCARP simulator*, and a framework for describing *in silico* experiments, referred to as *carputils* (Fig. 2). The *openCARP* simulator has been developed with the following objectives in mind:

- **Accessibility**: The main distribution mechanism of *openCARP* emphasizes binary packaging for commonly used platforms (currently Linux and macOS), platform-independent Docker containers, and a detailed documentation supporting source installations on all levels of computers including large-scale national HPCs. Only providing a source installation excludes major parts of the user community who are not sufficiently versed in compiling scientific software, or do not want to manage a complex software stack, especially given the release requirements and potential conflicts with compilers and underlying software libraries. The source code of all *openCARP* components is maintained in the public GitLab instance git.opencarp.org. *carputils* is published under the Apache 2.0 open source license, and the *openCARP* simulator under the [Academic Public License](https://opencarp.org/download/license) restricting the open use to academic non-profit cases. A commercial license can be requested.
- **Interoperability:***openCARP* builds on two decades of CEP modeling experience gained with its proprietary predecessors, *CARP* [16] and *acCELLerate* [17], with a stable user base about 130 registered users. The predecessor software packages have been extensively used by several CEP modeling groups and led to >250 peer-reviewed journal publications, covering the full range of modeling issues including numerical methods [71], anatomical modeling and model functionalization [48, 72] as well as a broad range of applications such as formation and maintenance of arrhythmias [73,74], EP therapies [75,76], diagnostic [77, 78] and therapeutic applications [79, 80]. The *openCARP* simulator has been built from scratch in terms of code, but the user interface is consistent with previous proprietary versions of *CARP* [16] to facilitate re-use of a large number of existing experiments. Moreover, *openCARP* interoperates with relevant community standards by providing input and output interfaces VTK [70] and *CellML* [51], for example, either directly or via other tools. All *openCARP* specific formats and standards are openly available and concisely described, with adequate tools for their management provided as source code.
- **Performance:***openCARP* has been implemented from scratch in C++ (2011 standard) but follows the same discretization and solver schemes that have been used successfully in the predecessor code *CARP* [16, 71]. Briefly, the *openCARP* simulator spatially discretizes the partial differential equations underlying the mono- or bidomain model (or potentially other physics like mechanics) using the FEM [81, 82] with linear basis functions. The bidomain equations are cast in the elliptic-parabolic form and decoupled to be solved sequentially [83], with various time stepping options including fully explicit Euler or *θ*-schemes, with or without operator splitting, leading to fully explicit or implicit-explicit solver schemes as the reaction term is always treated explicitly. The interested reader is referred to the openCARP user manual for numerical details. The FEM implementation makes use of parallel mesh management and mesh partitioning [68]. The resulting linear systems of equations are solved with *PETSc* [84], with various pre-configured solver options [85–87], under exploitation of parallel algebraic matrix-vector operations, parallel I/O, and timing routines among others. On each node, the embarassingly parallel *LIMPET* library [88] is used to calculate the cellular electrophysiology models. Time integration schemes for these ODE systems can be controlled per variable and include Runge-Kutta, Rush-Larsen, Rosenbrock [89] and advanced schemes accessible via *CVODE* [90] with adaptive step size and error control. While profiling and benchmarking plays a central role during development, *openCARP* does not yet employ detailed benchmarking or profiling during integration testing. Currently, integration tests are binned into different execution time groups and tests changing group are marked as failed.
- **Transparency:** The availability as source code will benefit the technically affine model developers in the CEP modeling community by allowing implementation of additional features, the critical revision of numerical schemes, and the identification of weaknesses. This will trigger constant re-engineering of the software and, thus, help improve software quality. The software is version controlled using git and connected to continuous integration / continuous delivery (CI/CD) services with integration testing of most simulation setups (currently 74 individual simulations). We have opted against unit testing at the current stage of development. Release versions of *openCARP* [91] are automatically archived and uploaded to the RADAR4KIT repository as part of the continuous deployment pipelines [92]. Submission information packages are built in BagIt format and comprise the source code, the binary versions for the support platforms, the respective version of the documentation and test reports. Metadata is automatically extracted from machine-readable files in the git repositories and provided according to the DataCite 4.3 scheme [93]. A DOI is minted for all published and archived versions to ensure long-term accessibility.

**Figure 2:**
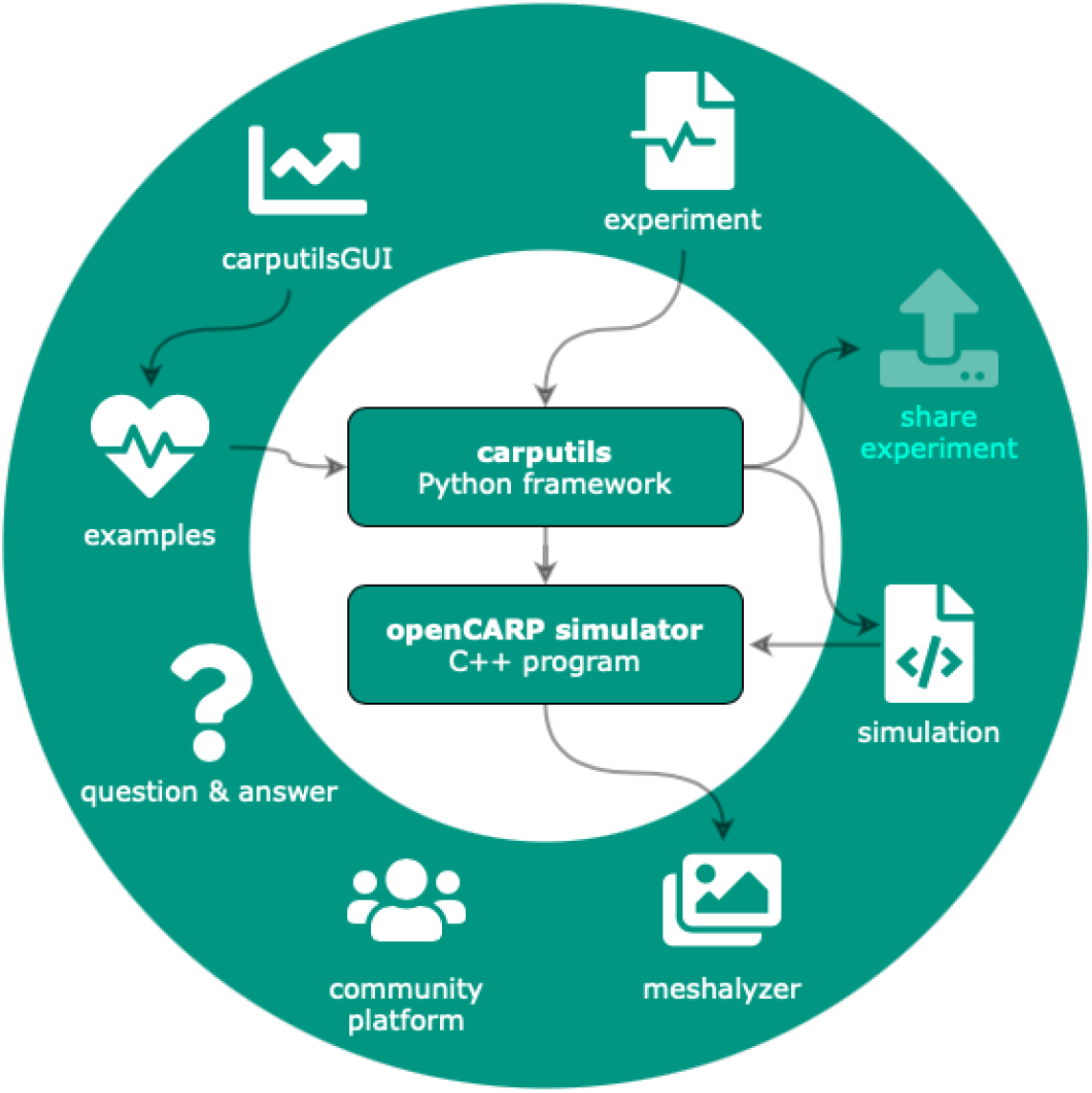
The *openCARP* ecosystem comprises several components that interact with its central pillars: *carputils* and the *openCARP* simulator. Users can write their own experiments using the *carputils* Python framework, or start with existing *examples. carputils* will set up the simulation, launch the *openCARP* simulator, perform postprocessing steps and call meshalyzer for visualization. Apart from the commandline interface, *examples* can also be parametrized, run and visualized using the web-based carputilsGUI. A community platform including a question & answer system completes the *openCARP* ecosystem.

*carputils* is a framework designed to define and execute *in silico* experiments by encoding the complex workflows of advanced CEP simulation studies in a reasonably standardized manner. By providing an additional abstraction layer on top of the *openCARP* simulator itself, we aim to achieve the following objectives:

- **Usability:** The main focus of *carputils* is on providing a modeling and simulation environment which enables researchers to carry out a wide range of studies out of the box. Only a physiological understanding of the problem at hand should be necessary for executing such experiments, not knowledge of underlying numerical details.
- **Reducing complexity:** *In silico* experiments defined with *carputils* facilitate learning by exposing only a small number of relevant parameters which are needed for controlling a given experiment.
- **Reproducibilty:** By providing mechanisms for easy sharing of *in silico* experiments, *carputils* contributes to the concept of Open Science and facilitates reproducibility and reuse. Owing to the complexity of experiments, the ability to share them has been severely hampered, even within the same laboratory, let alone across different ones. We overcome barriers posed by numerous differences in local installations, inconsistencies between code revisions, and the management of high dimensional input parameter spaces defining a setup: meshes, label fields, electrode definitions and pacing protocols, limit cycle state vectors, structural entities such as fibrotic lesions or scars, functional heterogeneities, conduction system topology, and parabolic solver method, to name but a few important ones.
- **Productivity:** *carputils* aims to boost productivity by providing standardized workflows for common tasks such as setting conduction velocities, computing limit cycles of ionic models to generate initial states, interrogate restitution properties, or post-processing and visualization of outputs.
- **Abstraction:** *carputils* uncouples technicalities of executing simulations on different platforms from experiment design such that they can be executed on any supported platform – from laptops to national HPC facilities – using the exact same command line. Platform-specific aspects related to scheduling and launching of jobs are hidden from the user, and encapsulated in abstract platform descriptions. Only generic user choices such as the number of processes to be used in a simulation must be provided.
- **Quality assurance:** Support for **regression testing** of all major simulator features is built in, including automated nightly building and testing of the entire software stack. *carputils* encodes a range of **verification benchmarks** such as the CEP N-version benchmark [9] and a suite of additional performance benchmarks of varying level of complexity, ranging from biventricular slices up to whole heart models, for measuring simulator performance, or for investigating the effect of numerical settings.

Additional open source software components have been developed to support the modeling, simulation and visualization when using *openCARP* as shown in Fig. 2. *meshtool* [54] (License: GPL v3) allows generating or interacting with geometric data.*[carputilsGUI](https://opencarp.org/documentation/carputilsgui)* (License: Apache 2.0) is a web-based graphical user interface to control specific *carputils* experiments and visualize results within the browser for low-threshold entry to *openCARP. meshalyzer* (License: GPL v3) is a tailored visualization program for CEP, even though VTK export allows seamless visualization in *[ParaView](https://www.paraview.org/)* [94] as well.

*openCARP* provides documentation for users with different levels of experience. [Video tutorials](https://opencarp.org/documentation/video-tutorials) introduce users to the basics of CEP modelling and guide them through their first steps with *openCARP*. Further videos introduce standard simulation pipelines and progressively advanced features covering the whole gamut from single cell simulations to 3D heart models including pre- and post-processing steps using the *carputils* framework. *carputils* [examples](https://opencarp.org/documentation/examples) cover common simulation scenarios and provide a starting point to develop custom *carputils* experiments in combination with the [API documentation](https://opencarp.org/documentation/carputils-documentation#master). The *openCARP* [reference documentation](https://opencarp.org/documentation/opencarp-parameters#master) contains all *openCARP* parameters to give full control of all aspects. Continuous deployment pipelines ensure that documentation (web page, PDF) is always in sync with the code by automatically generating these artifacts based on the code.

### 2.1 Workflows and Use Cases

In the following sections, we introduce the *openCARP* simulation framework by covering common workflows for use cases ranging from the cellular to the organ level. Specifically, we elucidate the key processing steps to implement a human biatrial model of persistent atrial fibrillation. An overview of a standardized processing workflow is given in Fig. 3.

**Figure 3:**
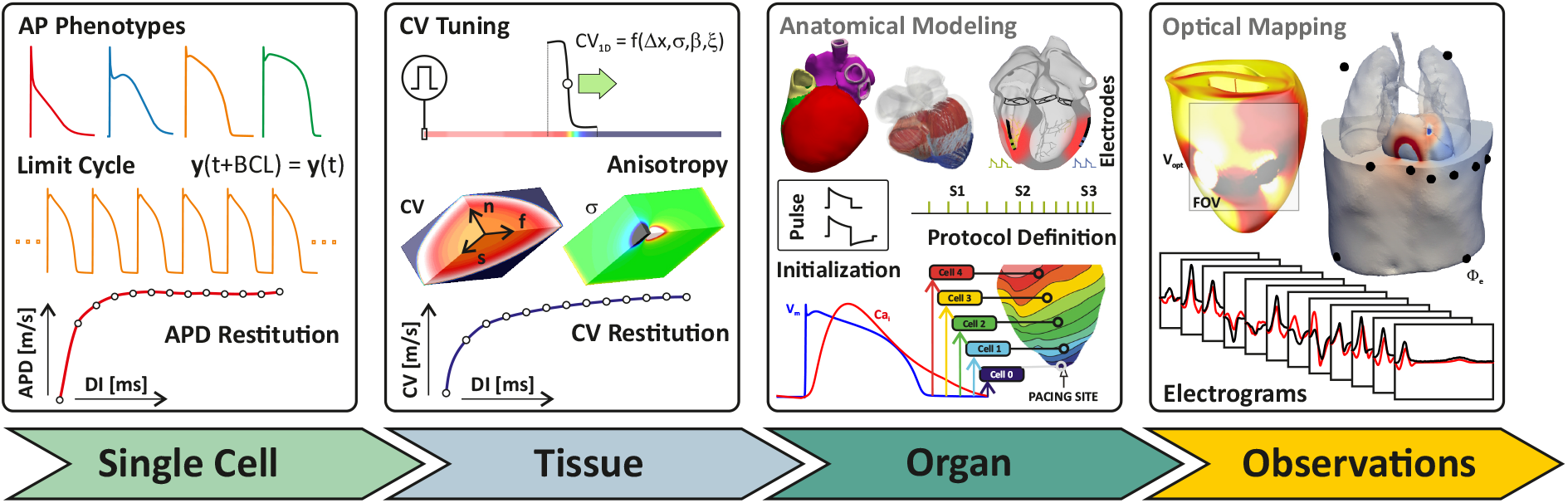
Overview of typical steps in an advanced CEP simulation study. Single Cell: AP phenotypes of different types of tissues are defined, paced to a stable limit cycle for a given set of cycle lengths, and dynamic properties such as APD restitution are evaluated. Initial state vectors ***y***_0_ are stored for each phenotype. Tissue: Conductivities *σ_ζ_* are determined for the mean spatial resolution Δ*x* of the organ-scale mesh, given surface-to-volume ratio, *β*, and numerical settings,, by simulating uniform wavefront propagation in 1D along an eigenaxis, *ζ*, to obtain the desired CVs along fiber, ***f***, sheet, *s*, and sheet-normal, ***n***, direction, and anisotropy ratios between intra- and extracellular domain. CV restitution is evaluated then by measuring CV under increasingly shorter diastolic intervals. Organ: Organ models are generated using anatomical modeling pipelines, currently not included in *openCARP*, consisting of mesh generation, definition of fiber architecture, labeling of regions, computation of anatomical reference frames [95–97] and possibly the incorporation of a cardiac conduction system. Location and geometry of electrodes is defined, as are the pulse shapes used for stimulation and the pacing protocol. The organ model is then populated by assigning the ionic model initial states ***y***_0_, as computed for each phenotype, to the respective tissues, and conductivities are assigned to the various regions to control CV. A pre-pacing protocol is applied to ascertain that the organ-scale model is as close as possible to a limit cycle for the given cycle length. Observations: Additional model components are set up to simulate observable data such as optical transmembrane voltages, *V*_opt_, invasively recorded electrograms **Φ**_e_, and non-invasive ECGs recorded from the body surface.

#### 2.1.1 Cellular Level

The ordinary differential equation (ODE) models representing cellular dynamics and ion current across the membrane are integrated in the *LIMPET* library. *openCARP* is shipped with a set of commonly used models (including Courtemanche et al. [98] Koivumäki et al. [99], ten Tusscher et al. [100], O’Hara et al. [101]). The commandline tool *bench* is an interface for carrying out single cell experiments to tune models of cellular dynamics to given (patho-) physiological conditions. The most basic use case is to compute APs for a given membrane model. Advanced features of *bench* are to control stimulation, compute restitution curves under various protocols, to carry out voltage clamp experiments or to clamp arbitrary other state variables.

~~~
bench --imp=Courtemanche --clamp-SVs=Ca_i --SV-clamp-files=Cai.dat
~~~

for example would clamp the intracellular calcium concentration of the Courtemanche et al. model to the time course defined in the `Cai.dat` file. The output files can either be processed with custom scripts or, ideally, pre- and postprocessing is integrated into a *carputils* Python script encoding the entire experiment including the call to *bench*. Besides CEP models, *bench* also provides myofilament models of active tension development. Often, one wants to change parameters of the cellular model to investigate how changes to a model component, due to e.g. drug effects, genetic mutations or disease-induced remodeling, affects behaviour. Parameters can easily be adjusted in *LIMPET* by specifying a value directly, or by using mathematical operations to alter the value relative to the default.

~~~
bench --imp=COURTEMANCHE
 --imp-par=“Gto-65%,GK1*2,GKs*2,GKur-50%,GCaL-55%,maxI_NaCa+60%,maxCa_up*0.5,C_m+20%”
~~~

for example would adjust the Courtemanche et al. model to reflect conditions of persistent atrial fibrillation-induced remodeling of cellular CEP [102] (Figure 4). The models equations are encoded in *[EasyML](https://opencarp.org/documentation/examples/01_ep_single_cell/05_easyml)*, a markup language developed for human readability. Changes to the model structure or parameters that are not exposed by default can easily be implemented in the `.model` files directly. An automated way to produce shared libraries that can be loaded at run time allows using these custom-built models or adapt numerical time integration without having to recompile the entire simulator. Additional models can be included using *CellML*, the XML-based community standard for ionic models. Models downloaded from the *CellML* repository [51] can be converted to *EasyML* using [tailored commandline scripts](https://opencarp.org/documentation/examples/01_ep_single_cell/10_fromcellml) or interactively via the third-party tool *[Myokit](https://www.myokit.org/)* [103].

**Figure 4:**
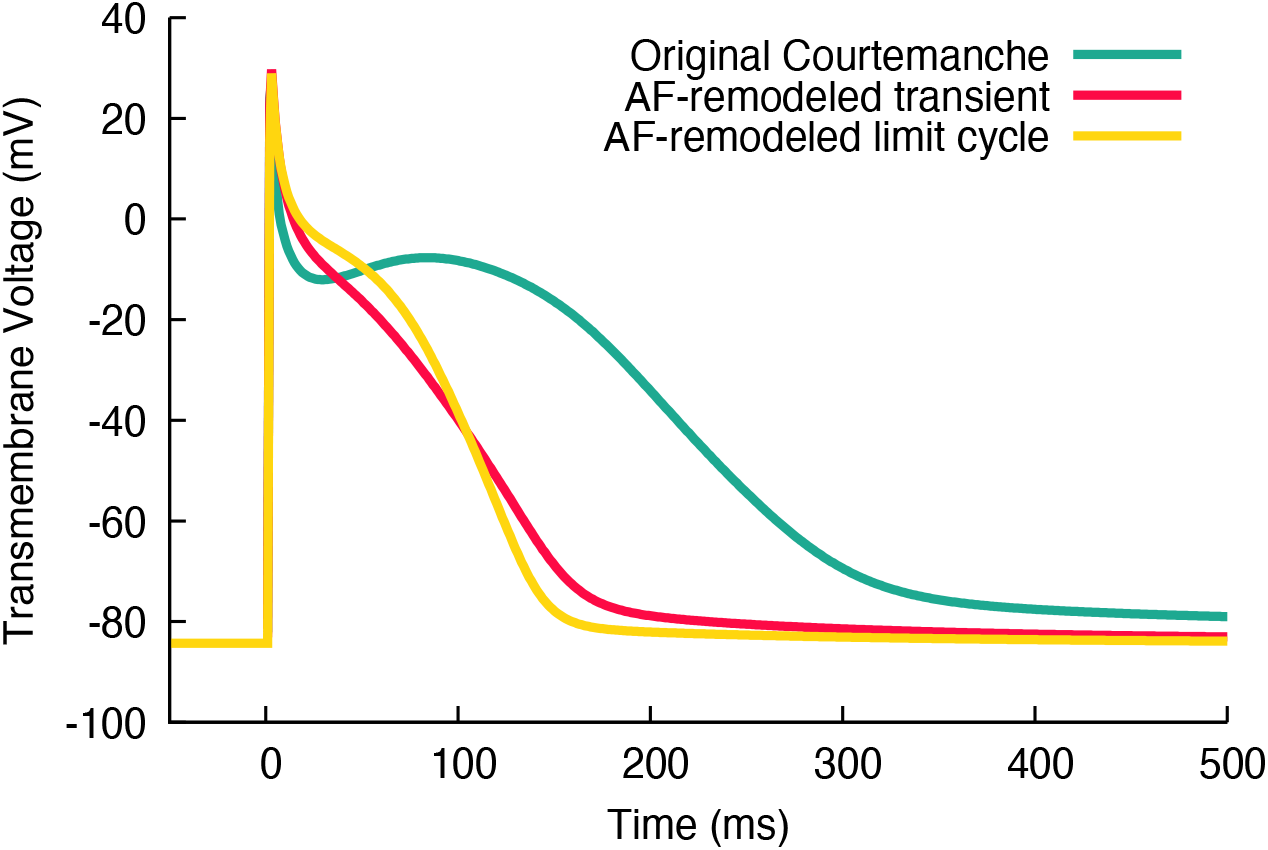
Action potentials of the original Courtemanche et al. [98] model (green) and remodelled variants reflecting persistent atrial fibrillation (AF) conditions [102]. Red: first action potential with the adapted parameters and initial state of the original model. Yellow: limit cycle AP after transient changes of the diastolic state equilibrated (100 stimuli at a basic cycle length of 1 s).

Initial conditions for tissue scale simulations are generated with *bench* by creating a set of myocyte models representing the CEP heterogeneity at the tissue scale. Each myocyte model is paced at one or more basic cycle lengths based on the activation rates desired for the tissue model until the model settles in to a limit cycle. Initial state vectors of each myocyte model are stored, to be used later to populate the tissue and organ scale model as detailed in the [limit cycle initialization](https://opencarp.org/documentation/examples/02_ep_tissue/03b_study_prep_init) *example*. Tailored experiments for interrogating and tuning of dynamic properties of ionic models such as [APD restitution](https://opencarp.org/documentation/examples/01_ep_single_cell/02b_apd_restitution) are also provided.

#### 2.1.2 Simple Tissue Level

Simple geometries allow investigating fundamental mechanisms and determining properties of tissue CEP. These simple geometries play an important role in the generation and interpretation of modeling results as one can investigate basic mechanisms by isolating the effects of confounding factors, such as the prevailing myocyte orientation (“fiber orientation”) or geometric complexities. Additionally, tissue behavioural properties can be quantified easily from these simplified models.

Mesh resolution has to strike a balance between computational effort and numerical accuracy. We refer to the N-version monodomain benchmark for spatial convergence considerations [9]. Tissue and cellular level properties need tuning to replicate activation and repolarization sequences as observed in wet lab experiments or the clinic. Stimulation patterns initiate propagation with a velocity which varies as a function of space. The anisotropic CV is determined by myocyte orientation, the tissue conductivities along and across the myocytes, the surface-to-volume ratio, and the fast sodium current of the cells, but modified by the curvature of the activation wavefront, and the refractory state of the sodium channels. Technical factors related to spatial discretization and time integration also play a role, particularly when using coarser spatial resolutions [9]. Specifically, the tissue conductivities need to be determined to ensure physiological direction-dependent CVs before running more complex simulations. This is easily performed using the *carputils* tool *tuneCV* [104]. A wave front propagation in a 1D strand is simulated using the desired AP phenotype, and the CV measured at the center of the strand for given numerical settings and spatial resolution. Thus, *tuneCV* compensates for artificial alterations in CV due to grid spacing, which is chosen to balance numerical accuracy and computational cost [9]. A comprehensive introduction to *tuneCV* can be found in the [*carputils example* “Tuning Conduction Velocity”](https://opencarp.org/documentation/examples/02_ep_tissue/03a_study_prep_tunecv). Other tissue properties like effective refractory period (ERP), vulnerable window or alternans can also be easily investigated in simple 1D geometries, for example, to extract arrhythmia predictors [28]. Additionally, rate dependent changes (restitution) of parameters provide insight into tissue behavior at fibrillatory activation rates. CV restitution is for example investigated in the [*carputils example* “CV Restitution”](https://opencarp.org/documentation/examples/02_ep_tissue/03d_conduction_velocity_restitution).

Simple 2D geometries can be used, for example, to investigate the effects of spatially heterogeneous tissue properties on arrhythmia dynamics. Spatial AP heterogeneity plays an import role in CEP as key features such as excitability, AP morphology and duration vary throughout the heart. Numerous effects of clinical relevance cannot be explained under the assumption of homogeneous tissue properties. Important examples are naturally existing heterogeneities within the ventricle that are responsible for the concordant T-wave in the ECG [105] and spatial changes of tissue conductivity (e.g. due to fibrosis) or/and electrophysiological properties in atrial fibrillation. In CEP modeling, two different approaches exist to account for spatial heterogeneity: either distinct regions are identified in which properties are uniform, or functions of space are defined which describe parameter variation, with granularity only being limited by spatial resolution. Region-based definition is, in general, easier to manage, but generates abrupt changes across region boundaries which may lead behavioural artifacts. Function-based assignment is more flexible and facilitates smooth variations that avoids potentially artificial high gradients, but is more challenging to define, particularly for complex anatomical models that, typically, require auxiliary anatomical coordinate systems to impose a desired variation [95–97]. The [*carputils example* “EP Heterogeneity”](https://opencarp.org/documentation/examples/02_ep_tissue/05c_cellular_dynamics_heterogeneity) illustrated the use of region-based heterogeneities, with four regions in which parameters can be set differently. Fig. 5A shows the impact of region-based variation in excitability upon the activation pattern. A comparison between region and gradient based heterogeneity is shown in [another *carputils example*](https://opencarp.org/documentation/examples/02_ep_tissue/05a_regions_vs_gradients).

**Figure 5:**
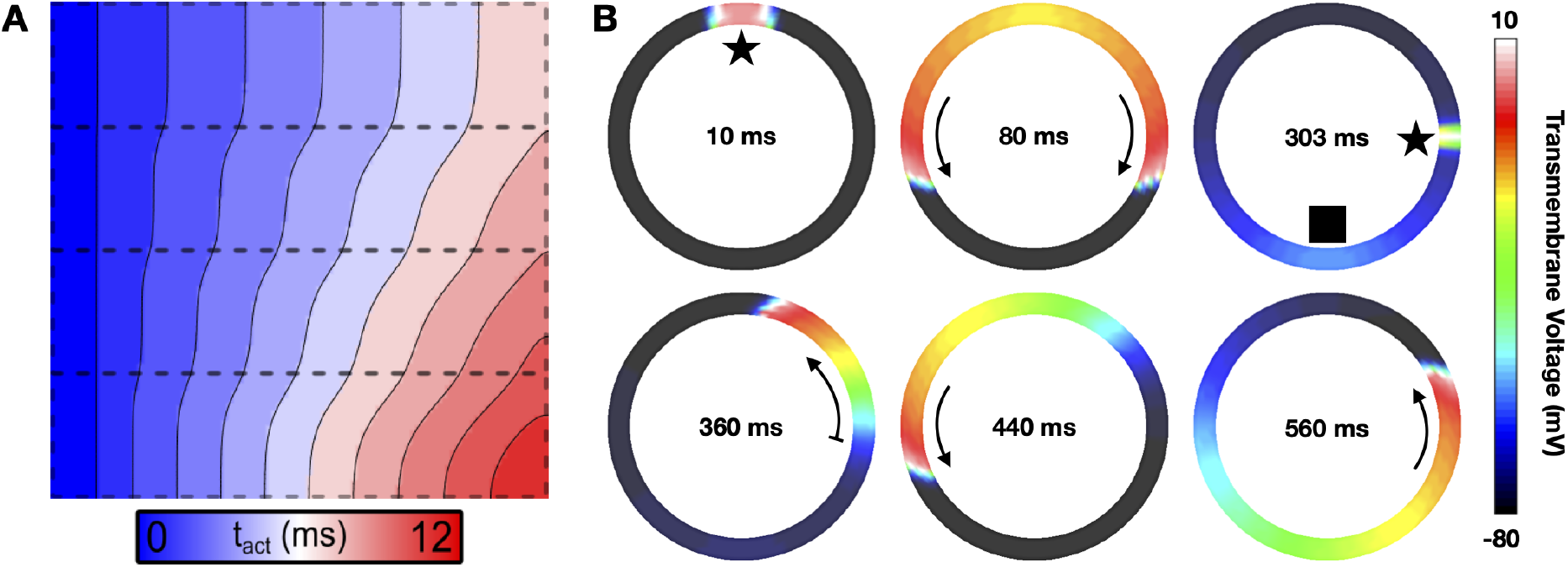
A: Activation times (*t_act_*) in a model with four regions of varying excitability. From the bottom to the top region, the conductance of the fast sodium channel (*g_Na_*) is set to 1x, 2x, 4x, and 8x *g_Na_* of the nominal value, respectively. Planar wave fronts initiated along the left edge of the sheet gradually distort due to these differences in CV. B: Initiation of a rotating wave in a ring model in a stylized representation of an atrial slice. The initial stimulus (delivered at 10 ms, marked by a star) induces bidirectional propagation (80 ms). At the end of the refractory period (303 ms), a second stimulus delivered at a different site located at a critical recovery isoline initiates a wave front that is blocked from propagating downwards where tissue is still refractory (rectangle). This leads to unidirectional propagation upwards (360 ms), setting up a sustained anatomical reentrant activation pattern (440 ms and 560 ms). This simulation was performed with ionic model settings corresponding to the persistent atrial fibrillation variant of the Courtemanche et al. model referred to above. In a healthy model, reentry cannot persist as the wavelength is too large for the given ring dimension.

Key advantages of *in silico* CEP models over *in vivo, ex vivo*, or *in vitro* models are their ability to precisely control all parameters, to observe all quantities of interest at high spatiotemporal resolution, to test large parameter spaces in a reproducible fashion, without being restricted by ethical concerns. Thus, mechanisms underlying CEP phenomena such as the formation and maintenance of an arrhythmia can be dissected in detail under a wide range of experimental settings. While organ-scale models are most comprehensive as they represent all factors at play, simplified 2D models are also highly relevant as these can provide crucial mechanistic insights with fewer confounding factors. Owing to their lighter weight, large parameter spaces can be probed and the observation of key variables and phenomena is more easily achieved than with the larger organ-scale models. Ring models representing a slice across a single cardiac chamber are a simple and popular 2D geometry for studying macro-reentrant arrhythmias. Fig. 5B illustrates the induction process of a reentrant wave in such a model. Reentrant activation is only sustained if the wavelength, defined as CV × ERP, is shorter than the perimeter of the ring. 2D sheet models can also be used to trace functional reentries, but their induction is less simple compared to the ring. The [*carputils example* “Induction Protocols”](https://opencarp.org/documentation/examples/02_ep_tissue/21_induction_protocols) presents four different methods to induce reentry in a sheet.

#### 2.1.3 Organ Level

For whole chamber or organ level models, meshes must be created that accurately reflect the anatomy of the entity to be modelled. Streamlined workflows for this task are currently under development by various labs [106], typically relying on connecting heterogeneous software components, including commercial software, that are specialized for particular processing stages, ranging from multi-label segmentation, meshing, fiber architecture generation [107], anatomical reference frame generation [95–97] or topology generation of the specialized cardiac conduction system [108]. The Python-based *carputils* is perfectly suitable for creating such workflows [109]. However, owing to the technological heterogeneity of all this software, a complete integrated workflow is currently not available within *carputils*, but many basic elements required for building such workflows are. High quality anatomical models equipped with realistic fiber architecture, multiple label fields and pre-computed anatomical reference frames are becoming publicly available. A point in case is the recent study by Strocchi et al. [35] that placed in a repository a cohort of 24 volumetric four-chamber meshes derived from heart failure patients, which can be used for organ level simulations with *openCARP*. For cohort studies in healthy individuals, statistical shape models allow covering even more anatomical variability. Biventricular and biatrial shape models, together with 100 volumetric instances ready to be used for *openCARP* simulation, are publicly available [110, 111]. It is anticipated and foreseeable that the number of publicly available anatomical twin models will further increase, hopefully in part due to the standardization of formats under *openCARP*, yielding an abundant pool of cardiac anatomy models available to *in silico* CEP research. For creating digital twin models based on data acquired from individual patients model parameters need to be identified that minimize the misfit between simulated and observed quantities [112]. Owing to their mechanistic nature, such digital twin models show promise as an approach towards realizing precision medicine where therapeutic responses simulated *in silico* may be used to guide therapy planning and delivery. For this purpose, largely automated pipelines have been proposed for the atria [72, 113], the ventricles and the whole heart [35,53]. The geometric models can be augmented with population-level *a priori* knowledge regarding myocyte orientation in the atria [114, 115], the ventricles [107] or the whole heart [116]. Like anatomy, functional properties can be represented by population averages [117] or personalized using non-invasive or intracardiac measurements [72, 79, 118–121]. Depending on the scientific question, relevant parameters can be global or spatial distributions, and can include, for example, tissue conductivity, ionic model properties, or the initially activated sites. Generic frames of reference, such as the universal ventricular coordinates [95] or universal atrial coordinates [122] allow parametrizing locations within the cardiac chambers to facilitate optimization approaches and transferability between models.

A common use case for organ level simulations is assessing the effects on arrhythmia inducibility of factors like fibrosis, scar or drugs. *carputils* provides several arrhythmia induction methods [123] to efficiently build such *in silico* experiments. The basic use is illustrated in [one of the currently 28 *carputils examples*](http://opencarp.org/documentation/examples/02_ep_tissue/21_induction_protocols) in addition to the API documentation. Users can readily integrate these induction methods in their own *carputils* experiments.

~~~
model.induceReentry.PEERP(…, stim_points, …)
~~~

calls the *carputils* function *PEERP* to apply a protocol which paces at the end of ERP (PEERP) from all points included in the `stim_points` list. Depending on the cellular and tissue properties, reentry can be induced as shown in Fig. 6A where a rotor at the left atrial appendage drives reentry in both atria. *Meshalyzer*, part of the *openCARP* ecosystem, was used to produce Fig. 6A. Alternatively, *openCARP* results can be directly output in VTK format, or converted to VTK format in a post-processing step, for seamless visualization in *ParaView* [94].

**Figure 6:**
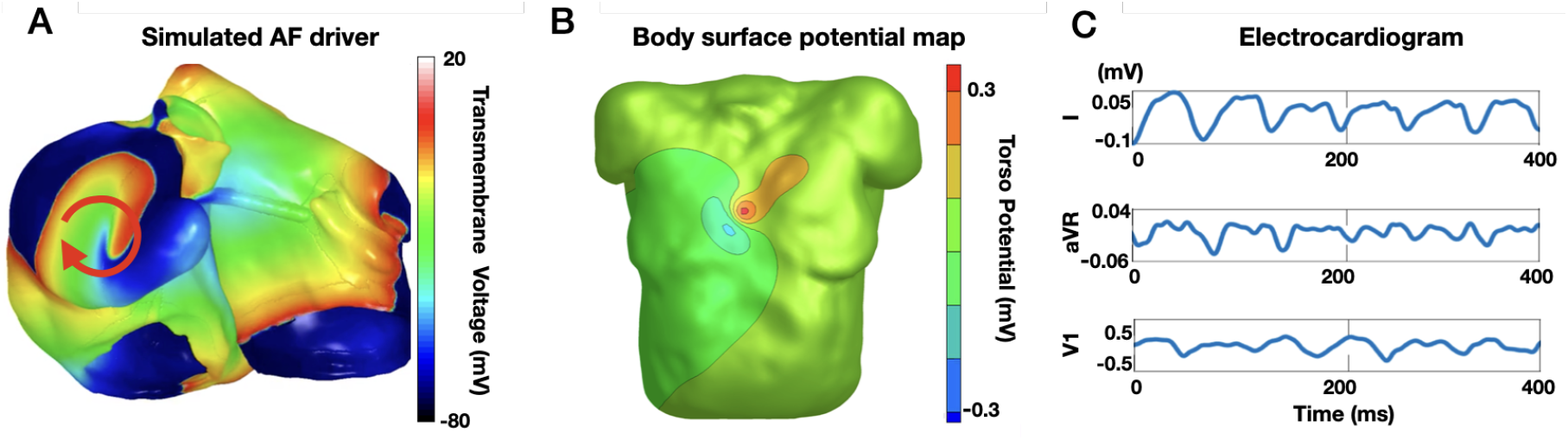
A: Biatrial volumetric model with ongoing reentry induced by pacing at the end of the refractory period using the *carputils* function `model.induceReentry.PEERP()`. B: Forward-calculated body surface potentials stemming from the transmembrane distribution in panel A. C: ECG (Einthoven lead I, Goldberger lead aVR and Wilson lead V1) extracted from the body surface potential map time series (panel B) reflecting the atrial fibrillation perpetuating at the tissue level (panel A).

#### 2.1.4 Body Level

Cardiac excitation and the resulting spatially heterogeneous transmembrane voltage distribution, *V*_m_(***x**, t*), in cardiac tissue act as sources that generate electric current flow in the extracellular domain comprising the interstitial space as well as a volume conductor potentially surrounding the cardiac tissue (blood pool, torso), often referred to as bath. Currents in the volume conductor set up an extracellular potential field, **Φ**_e_(***x**, t*), that can be sampled by electrodes to acquire body surface potentials or intracardiac electrograms. Apart from experimental settings where cardiac sources in the form of *V*_m_(***x**, t*) can be observed more directly [124, 125], extracellular potential recordings are the only means in the clinic to infer the cardiac source distribution. Owing to the unique importance of these signals for diagnosis and guiding therapy, methodologies facilitating their accurate simulation are an indispensable adjunct in many CEP simulation studies. These demands are reflected in the underlying bidomain equations solved by *openCARP*, which comprise both computational domains, the intracellular domain and the extracellular domain, thus facilitating the simulation of electrograms [75, 76, 126] or torso potentials [78, 120]. The electrical potential within the volume of interest can be computed in *openCARP* using different methods depending on the trade-off between computation and accuracy desired: i) The bidomain setting, which considers bath-loading effects [127] and allows for bidirectional interplay between intracellular and extracellular potential fields; ii) Alternatively, extracellular potential can be recovered in a post-processing step from a given spatiotemporal transmembrane voltage distribution obtained through, a monodomain simulation. Bishop & Plank [128] showed that the latter approach (referred to as *pseudo bidomain*) retains accuracy in most cases, particularly with augmentation techniques to recover bath-loading effects close to the tissue-bath interface [129], while drastically reducing the computational cost; and iii) An even more simplistic approach is to assume that the cardiac tissue is immersed in a homogeneous volume conductor of infinite size, which allows recovering the extracellular potential **Φ**_e_ at specific points in space from a given transmembrane voltage distribution by using the integral solution of Poisson’s equation. [One of the *carputils examples*](https://opencarp.org/documentation/examples/02_ep_tissue/07_extracellular) introduces all three methods and involves the user in simple experiments highlighting the differences between them. To compute surface potentials in own *carputils* experiments, the `ep` module provides the `model_type_opts(sourceModel)` function to choose between **Φ**_e_ recovery, pseudo-bidomain and bidomain. Fig. 6B shows a snapshot of the body surface potential map generated by the transmembrane voltage distribution *V*_m_ during atrial fibrillation shown in Fig. 6A. By extracting the time series of extracellular potentials **Φ**_e_(*t*) at the electrode locations, virtual ECGs can be derived by subtraction of electrograms along the lead axes (Fig. 6C).

## 3 Discussion

*openCARP* is a CEP simulation environment for carrying out advanced *in silico* experiments. The *openCARP* framework is highly versatile and comprehensive, in principle allowing replicating and building on the vast majority of published CEP simulation studies which rely only on the monodomain, bidomain and Poisson equations. *openCARP* builds on the core technologies of its proprietary predecessors, *CARP* and *acCELLerate*, that have matured over 20+ years of cutting edge modeling research, having been used in more than 250 published studies. *openCARP* provides a convenient and flexible user interface that enables performing complex simulations, requiring little to no programming experience. It supports the full research life cycle from exploration through to conclusive analysis and publication, to archiving and sharing of data, experimental protocols, models and source code. As such, *openCARP* can be a suitable software solution for a large portion of the CEP community, by contributing to the use, transparency, standardization and reproducibility of *in silico* approaches.

*openCARP* is designed to appeal to users of all experience levels. New users can easily explore parameter effects in prebuilt *carputils* experiments, while more experienced users can further extend such experiments to more elaborate scenarios, or build their own complex experiments from scratch. *openCARP* provides a solution for potential new users in the fields of basic science CEP (integrating experimental data into mechanistic models), cardiology research (understanding arrhythmia mechanisms, diagnostic support, therapy stratification, planning and delivery), pharmaceutical companies (virtual testing arrhythmogenic potency of new compounds), medical device companies (interpreting measured signals, optimizing new device designs [130], testing their safety and efficacy, and understanding reasons for device failure [26]), educational use (visualizing complex heart function), and regulatory entities (e.g., the FDA which is already encouraging the use of numerical models for drug and device testing [2]).

The central resource and entry point is the project web page www.openCARP.org. From there, the community has access to the user and developer documentation, tutorials, *examples*, the GitLab projects, and a question and answer system [131]. Besides the online platform, in-person contact among and between users and developers is recognized as a key factor to build and maintain a strong community. Regular user meetings allow training new users, exchange experiences with the software, and stay close with the community as part of a continuous feedback loop. Within a year from the release of the first public version, more than 250 users opened an account in our user and development community.

*openCARP* is a meritocratic, consensus-based community project. Anyone with an interest in the project can join the community, contribute to the project design, and participate in the decision making process. Community contributions are highly appreciated and can range from code commits to bug reports, suggestion of ideas and new directions for improving the software, documentation or any aspect of the community platform. Code contributions will enter a review process as a quality control measure and will be integrated in the code base if they meet quality criteria and satisfy a community need. The roles in the *openCARP* project are *users*, *developers*, *maintainers* and the *steering committee*. Users are community members who have a need for the project. The *openCARP* project asks its users to participate in the project and community as much as possible. Contributors engage with the project in concrete ways, through the issue tracker, question and answer system, or by writing or editing documentation. They submit changes to the project itself via commits to non-protected branches of our git repositories, which then undergo review by the maintainers before be merged into the main branch of code. Most of the 12 maintainers and five steering committee members are employed on tenured stable academic positions, ensuring long-term maintenance and support.

An important prerequisite for the reproducibility and reusability of models and associated simulation data is their enrichment with metadata, and publication as FAIR [132] and open data. While *openCARP* carries the features to reproduce the majority of published CEP simulations, the practical limitation is a not fully complete description of experiments or limited data availability. We are, therefore, currently embedding mechanisms in *carputils* to automate the extraction of all relevant information for a specific experiment, and create packages suitable for archiving and sharing. A convenient interface to upload these simulation packages to an *openCARP* specific section of the [RADAR4KIT research data repository](https://radar.kit.edu/) [133] will be provided in the near future. RADAR4KIT is a long-term research data repository that adheres to the Open Archival Information System (OAIS) standard [134]. It provides access control (public, private sharing via link), persistent DOI identifiers, standardized interfaces facilitating data harvesting, and is integrated with research data meta repositories like re3data [135]. In this way, the *carputils* experiment can be further evaluated by the scientific community, and future research can build upon it. Errors or previously undiscovered artifacts can be found and described, leading to improved data and research quality in the long term as one of the main goals of open science.

In conclusion, *openCARP* is an open CEP simulator released to the academic community to advance the computational CEP field by making state-of-the-art simulations accessible. A commercial license can be requested. In combination with the open source *carputils* framework and additional open source software around it, it offers a tailored software solution for the scientific community in the CEP field and contributes towards increasing use, transparency, standardization and reproducibility of *in silico* experiments.

## Acknowledgments

We gratefully acknowledge support by Deutsche Forschungsgemeinschaft (DFG) (project ID 391128822, LO 2093/1-1, GS 1758/4-1). We further acknowledge the software contributions of NumeriCor GmbH, Graz, Austria who conceived and implemented the *openCARP* simulation core. We thank Felix Bach, Jochen Klar and Robert Ulrich for developing and providing the *[openCARP development environment](https://git.opencarp.org/openCARP/openCARP-CI)* as well as Michael Selzer and Philipp Zschumme for developing and providing *[toolcompendium](https://git.opencarp.org/iam-cms/toolcompendium)* for generation of user documentation. We thank Prof. Natalia Trayanova for agreeing to make the *EasyML* model generator, developed in her lab by Rob Blake, available to be distributed with *openCARP*. We thank Giorgio Luongo for providing figures of simulation results.

